# Rapid evolution of mammalian HERC6 and independent duplication of a chimeric HERC5/6 gene in rodents and bats suggest an overlooked role of HERCs in antiviral innate immunity

**DOI:** 10.1101/2020.09.14.294579

**Authors:** Stéphanie Jacquet, Dominique Pontier, Lucie Etienne

## Abstract

The antiviral innate immunity in mammals has evolved very rapidly in response to pathogen selective pressure. Studying the evolutionary diversification of mammalian antiviral defenses is of main importance to better understand our innate immune repertoire. The small HERC proteins are part of a multigene family including interferon-inducible antiviral effectors. Notably, HERC5 inhibits divergent viruses through the conjugation of ISG15 to diverse proteins-termed as ISGylation. Though HERC6 is the most closely-related protein of HERC5, it lacks the ISGylation function in humans. Interestingly, HERC6 is the main E3-ligase of ISG15 in mice, suggesting adaptive changes in HERC6 with implications in the innate immunity. Therefore, HERC5 and HERC6 have probably diversified through complex evolutionary history in mammals, and such characterization would require an extensive survey of mammalian evolution. Here, we performed mammalian-wide and lineage-specific phylogenetic and genomic analyses of HERC5 and HERC6. We used 83 orthologous sequences from bats, rodents, primates, artiodactyls and carnivores – the top five representative groups of mammalian evolution and the main hosts of viral diversity. We found that mammalian HERC5 has been under weak and differential positive selection in mammals, with only primate HERC5 showing evidences of pathogen-driven selection. In contrast, HERC6 has been under strong and recurrent adaptive evolution in mammals, suggesting past genetic arms-races with viral pathogens. Importantly, we found accelerated evolution in the HERC6 spacer domain, suggesting that it might be a pathogen-mammal interface, targeting a viral protein and/or being the target of virus antagonists. Finally, we identified a HERC5/6 chimeric gene that arose from independent duplication in rodent and bat lineages and encodes for a conserved HERC5 N-terminal domain and divergent HERC6 spacer and HECT domains. This duplicated chimeric gene highlights adaptations that potentially contribute to rodent and bat antiviral innate immunity. Altogether, we found major genetic innovations in mammalian HERC5 and HERC6. Our findings open new research avenues on the functions of HERC6 and HERC5/6 in mammals, and on their implication in antiviral innate immunity.

## Introduction

As a result of sustained exposure to viral infections, mammals have evolved a sophisticated and diversified immune repertoire against viruses. A hallmark of mammalian antiviral immunity is the induction of type I interferon (IFN) upon viral infection. This cytokine upregulates the transcription of hundreds of interferon-stimulated genes (ISGs) in viral infected cells (1). Many of these ISGs encode proteins with antiviral functions, named restriction factors, which are critical players in the first line of the innate immune defense inhibiting different steps of the viral replication cycle (2).

Viruses have adapted to circumvent, subvert, or antagonize these host restriction factors (3). Reciprocally, restriction factors have rapidly and repeatedly evolved to maintain defenses against evolving viral pathogens, leading to virus-host evolutionary arms-races (3,4). These dynamics of reciprocal adaptations can leave genetic signatures in the host restriction factors. Significant accumulations of non-synonymous changes over synonymous substitutions – designated as positive selection, as well as codon deletions or insertions that may alter the virus-host interface, are common genetic signatures of such long-term evolutionary arms-races (3–6). At the genomic level, gene duplication and recombination are among the most important mechanisms that can diversify the antiviral immune repertoire. In particular, gene duplication can generate adaptive molecular novelty allowing hosts to escape viral antagonism, evolve new immune functions, or increase the depth of antiviral response (3,7,8). The weight of such evolutionary mechanisms in mammalian immunity is highlighted by the extent of multigene families, which encode important ISG-encoded proteins, such as the Tripartite Motif-containing (TRIM) (9–12), Apolipoprotein B Editing Complex (APOBEC3) (13–15), Interferon-induced Protein with Tetratricopeptide Repeats (IFIT) (16–18), Interferon induced Transmembrane *protein* (IFITM) (19) families. For example, the APOBEC3 family has expanded in a lineage-specific manner in primates (20), artiodactyls (21) and bats (15), generating variability in mammalian antiviral response (14). However, the evolutionary and functional diversification of many antiviral families remains poorly characterized in mammals. Deciphering the evolutionary trajectories of multigene family members can provide insights into the genetic mechanisms underlying the diversification of antiviral responses and may allow identifying novel antiviral proteins.

The HECT and RLD domain containing E3-ubiquitin protein ligases, known as HERC proteins, are encoded by a multigene superfamily that is poorly studied in mammals. With six gene members, the HERC family is divided into two subfamilies, the large (HERC1 and HERC2) and the small (HERC3-6) HERC proteins (22). The small HERCs are structurally characterized by a N-terminal RCC1-like domain (RLD), a spacer region, and a C-terminal HECT (Homologous E6-AP Carboxyl Terminus) ubiquitin E3-ligase domain, while the large HERCs possess at least two RLD domains in addition to a HECT domain (23). This structural difference between large and small HERCs reflects their independent evolutionary history (24). In the antiviral immune context, much attention has been devoted to the small HERCs, in particular to HERC5 - an ISG-encoded antiviral effector - and HERC6 its closest relative (24,25). In humans, HERC5 acts as a HECT ubiquitin and E3-ligase (26–28). HERC5 notably conjugates the ubiquitin-like protein ISG15 to different protein targets, a process termed ISGylation (29–31). The protein targets may be non-specific newly synthetized viral proteins, specific viral proteins, or specific host proteins (29–31). ISGylated proteins are modified, functionally disrupted or altered in their localization within the cells. Through this ISGylation activity, HERC5 has an antiviral function against highly divergent viruses, including retrovirus (Human and Simian immunodeficiency viruses, HIV and SIV), papillomavirus, and influenza virus (25,31–33). For example, HERC5 targets the early stage of HIV assembly by catalyzing the ISGylation of the viral Gag protein (30), while it reduces influenza A viral replication through the conjugation of ISG15 to the viral NS1 protein (31). Besides, HERC5 appears to further interfere with HIV replication in an ISGylation-independent manner by impacting the nuclear export of Rev/RRE-dependent viral RNA, most likely through determinants in the RLD domain (33). In contrast, although HERC6 is the most closely-related protein of HERC5, little is known about its functional implication in mammalian antiviral immunity (25). The antiviral role of HERC6 has mainly been described in mouse, in which the HERC5 gene has been lost and functionally substituted by HERC6, the main murine E3-ligase of ISG15 (28,34,35). In humans, although the HERC6 protein possesses a HECT E3-ligase domain, it is devoid of ISGylation function (25).

These evolutionary and functional differences between mammals suggest lineage-specific adaptive changes in HERC5 and HERC6. Two previous studies showed that HERC5 and HERC6 genes have evolved under positive selection during vertebrate evolution (25,36). They further pointed to the RLD domain as the main target of the selective pressure. While these studies have provided important insights into the diversification of HERCs, the evolutionary analyses have certain limitations: (i) the scarcity of species analyzed (only 12 species, *versus* 81 species with at least 10 species per order in this current study), (ii) the overrepresentation of primates compared to other mammalian species (seven primates, two to three carnivores, two artiodactyls and one perissodactyl), (iii) the integration of highly divergent species, which may bias the genetic inferences by increasing the number of false positives (37). Moreover, a recent study in primates have shown differences in HERC5 and HERC6 selective pressures (38). Therefore, how HERC5 and HERC6 genes have evolved within mammalian orders has not been fully deciphered. Nor is the evolutionary dynamic of HERC5 and HERC6 expansions and contractions across mammals.

Here, we decipher the evolutionary history of mammalian HERC5 and HERC6 via mammalian-wide and lineage-specific phylogenetic and genomic analyses. We analyzed the orthologous sequences of HERC5 and HERC6 from bats, rodents, primates, artiodactyls and carnivores – the top five representative groups of mammalian evolution and the main hosts of viral diversity. First, we show that HERC6 - and to a much lesser extend HERC5 - has been under strong positive selection. Second, we stressed the HERC6 spacer region as a potential pathogen-mammal interface, targeting viral proteins and/or being the target of virus antagonists. Finally, we identified independent gene duplications through recombination between HERC5 and HERC6 in some bat and rodent lineages, which have led to the fixation of a chimeric HERC5/6 gene in both mammalian orders. Taken together, our results suggest that HERC6 may be an important antiviral protein in mammals and identified a novel chimeric HERC member in bats and rodents that may contribute to unique antiviral responses in these species.

## Materials and Methods

### Collection of mammalian HERC5 and HERC6 orthologous sequences

Full-length HERC5 and HERC6 coding sequences were analyzed in bats, rodents, primates, artiodactyls and carnivores - the top five mammalian orders in terms of zoonotic viral diversity they host (39,40). HERC5 and HERC6 coding sequences from each group were obtained using the Little Brown bat (*Myotis lucifigus*), mouse (*Mus musculus*), human (*Homo sapiens*), cow (*Bos taurus*) and dog (*Canis lupus familiaris*) Refseq proteins as queries, respectively, through tBLASTn searches of the “Nucleotide” database in GenBank (41,42). The species and accession numbers are presented in Supplementary Table1.

### Characterizing the evolution of HERC5 and HERC6 synteny in mammals

The genomic locus of HERC5 and HERC6 genes in Little Brown bat, mouse, human, dog and cow were obtained from Ensembl (www.ensembl.org/index.html), and GenBank Refseq genome database (42). Their coding sequences were used as queries for BLASTn searches against a total of 110 whole genome assemblies from bats, rodents, primates, carnivores and artiodactyls (Supplementary Table1) available in GenBank database (42). We analyzed eight additional mammalian genomes from Proboscidea, Lagomorpha, Scandentia, Perissodactyla, Sirenia, Soricomorpha, Eulipotyphla, and Tubulidentata orders (*Orycteropus afer, Loxodonta Africana, Trichechus manatus, Tupaia chinensis, Condylura cristata, Ceratotherium simum, Equus przewalskii* and *Sorex araneus*, respectively) (Supplementary Table1). The synteny of HERC5 and HERC6 genes was analyzed and visualized through BLAST searches against the 110 annotated genomes in GenBank database (42). Newly identified HERC-like paralogs (see results) were confirmed by blasting and aligning their whole sequence (intron and exon regions) to the genomic region containing HERC5 and HERC6 genes in three bat species (*Myotis lucifigus, Rhinolophus ferrumequinum* and *Phyllostomus discolor*), and three rodent species (*Chinchilla lanigera, Mastomys coucha, Cavia porcellus*), which are representative genomes with good assembling quality (based on the N50, the number of scaffolds, and sequencing coverage).

HERC5 and HERC6 orthologs as well as HERC-like paralogous sequences were aligned for each mammalian order separately using the program MACSE (43), and the alignments were manually curated. A phylogenetic tree was then built for each gene and mammalian order, and for a combined dataset of HERC5 and HERC6 genes (rooted with HERC3 as an outgroup, which is the most closely related gene to HERC5 and HERC6), using the maximum likelihood method implemented in the ATGC-PhyML Web server (44). Each phylogenetic tree was based on the best substitution model (GTR+G+I), as determined by the Smart Model Selection (SMS) program in PhyML (45) and node statistical support was computed through 1,000 bootstrap replicates.

### Assessing recombination events in HERC5 and HERC6 paralogs and orthologs

To test whether recombination has occurred in HERC5, HERC6 and HERC-like genes, we run the GARD (Genetic Algorithm for Recombination Detection) method (46) implemented in the HyPhy package (47,48), using a general discrete site-to-site rate variation with three rate classes. The program uses a genetic algorithm to screen multiple-sequence alignment for putative recombination breakpoints and provides the probability of support for each breakpoint. GARD analyses were run for each mammalian order and each gene separately, including the newly identified HERC-like paralog.

### Positive selection analyses of HERC5 and HERC6 coding sequences in mammals

To determine whether HERC5 and HERC6 have been targets of natural selection during mammalian evolution, we carried out positive selection analyses on orthologous coding sequences from bats (n=10 and 13, respectively), rodents (n=11 and 16), primates (n=19 and 20), carnivores (n=20 and 23) and artiodactyls (n=21 and 11). As combining highly divergent sequences for positive selection analyses can lead to misleading results, we performed positive selection analyses on separate dataset for each mammalian order. For Artiodactyls, three different datasets were analyzed: the first dataset included all the available species, the second was restricted to cetacean species, and the third excluded the cetaceans. Analyzing each mammalian order separately allowed us to qualitatively compare the evolutionary profile of both genes in each mammalian order. We first tested for positive selection at the gene level using two different methods available in the Codeml program, which is implemented in the PAML package (49). This program allows both gene- and site-specific detection of positive selection by comparing constrained models that disallow positive selection (models M1 and M7) to unconstrained models allowing for positive selection (M2 and M8). We ran the different models with the codon frequencies of F61 and F3×4 with a starting omega ω (*dN/dS* ratio) of 0.4. Likelihood ratio tests were computed to compare the models (M1 vs M2 and M7 vs M8), and codons evolving under significant positive selection (*dN/dS* >1) were identified using the Bayes Empirical Bayes (BEB) with a posterior probability ≥ 0.95. The residues under positive selection were further assessed using two other methods, the Fast-Unbiased Bayesian Approximation (FUBAR) (50) and the Mixed Effects Model of Evolution (MEME) (51), both implemented in the HYPHY package. To increase the specificity of our results, we only kept the sites that were significantly identified by at least two of the four methods used. When significant recombination breakpoints were detected, positive selection analyses were carried out for each fragment identified by GARD. Similarly, we sought for signatures of adaptive selection in the HERC-like paralogs of rodents (see Results), for which five orthologous coding sequences were available. Furthermore, we tested if the three HERC domains (RLD, spacer region and HECT) have similarly been subjects of positive selection, by analyzing each domain separately.

Finally, to determine if HERC5 and HERC6 have experienced episodic selection within mammalian orders, we carried out the branch-specific analysis aBSREL (52,53), implemented in the HYPHY package. This program allows testing the significance of positive selection and quantifying the *dN*/*dS* ratio for each branch independently.

## Results

### Lineage-specific changes in HERC5 and HERC6 copy number

To determine the genomic evolutionary history of HERC5 and HERC6 in mammals, we performed a complete synteny analysis for 14 chiropteran, 25 rodent, 25 primate, 23 artiodactyl, and 23 carnivore species, and eight more species from different orders. We first found that HERC5 and HERC6 synteny is mostly conserved throughout eutherian mammals (Figure 1, Supplementary Figure 1). However, we also detected that gene erosion has repeatedly shaped the evolution of HERC5 and HERC6 in mammals. While primate and carnivore genomes encode both genes, the artiodactyls and the rodents have experienced gene loss or pseudogenization, generating HERC gene copy number variation in each group. Indeed, although all the studied artiodactyl species encode an intact HERC5 gene, the cetacean genomes showed HERC6 pseudogenization, through nucleotide deletions that impacted frameshift, as well as early stop codons (Figure 1, Supplementary Figure 2). Moreover, we confirmed the erosion of HERC5 in rodents (54), and we further show that this loss has most probably occurred in the common ancestor of the Cricetidae and Muridae. Interestingly, we also did not find the HERC5 gene in the rhinoceros genome, suggesting at least two independent losses of HERC5 in mammals, specifically in the Rodent and Perissodactyl orders (Figure 1).

**Figure 1.**
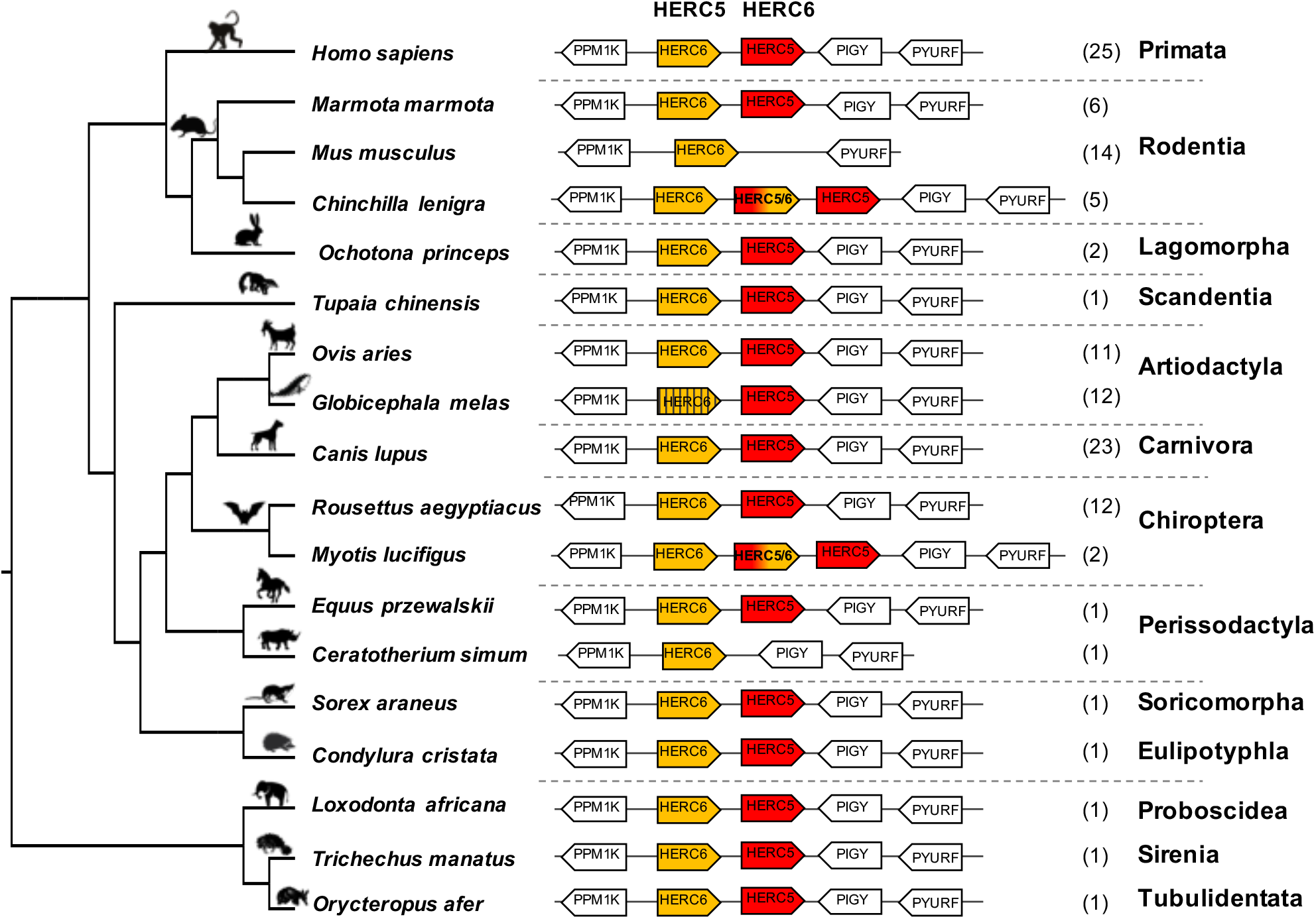
Evolutionary dynamics of mammalian HERC5 and HERC6 gene loci. Representation of the HERC5 and HERC6 gene loci from mammalian genomes. Plain colored arrows represent intact HERC5 and HERC6 genes, striped colored arrows indicate HERC5 or HERC6 pseudogenes, and white arrows are adjacent syntenic genes. The newly identified chimeric HERC5/6 genes are bicolored. The numbers in brackets indicate the total number of genomes analyzed which contain the corresponding genomic organization in each mammalian order. In primates and carnivores, the HERC5 and HERC6 genes are well conserved. In the cetacean and rhinoceros’ species, the HERC6 has been pseudogenized and lost, respectively, while rodent HERC5 has been lost in the Muridae and Cricetidae families. A duplicated HERC5/6 fused gene is independently fixed in several rodent species and in the *Myotis* genus in bats.

Finally, in addition to these independent losses of HERC5 or HERC6 in mammals, we found multiple evidences of HERC5 duplication followed by pseudogenization in primate (*Sapajus apella, Cebus capucinus, Aotus nancymaae and Saimiri boliviensis*), carnivore (*Canis lupus, Vulpes Vulpes, leptonychotes weddellii*) and artiodactyl (*Ovis aries*) species, highlighting a strong dynamic of gene gain and loss in mammalian HERC5 and HERC6.

### Ancient and recent recombinations have shaped the evolution of a duplicated chimeric HERC5/6 gene in rodents and bats

While most mammalian species possess one or both HERC genes, we found independent duplications of HERC5 in the chiropteran *Myotis* genus and the rodent Hystricognathi infra-order (Figure 1, Supplementary Figure 3). These duplicated genes were identified in three bat *Myotis* species (*M. lucifigus, M. brandtii*, and *M. davidii*) and five rodent species (*Fukomys damarensis, Heterocephalus glaber, Cavia porcellus, Chinchilla lanigera* and *Octodon degus*) (Table S1). This dates these duplication events to at least 30 MYA (million years ago) and 44 MYA, respectively.

Surprisingly, alignments of HERC5, HERC-like and HERC6 proteins in each mammalian group reveal 96 - 99% amino acid identity between the HERC5 and HERC-like N-terminals, and 74 - 84% amino acid identity between the HERC-like and HERC6 C-terminals (Figure 2a, Supplementary Figure 4). To test whether this may be reminiscent of recombination in rodents and bats, we used the GARD program. We identified a significant recombination breakpoint located upstream from the spacer region at 1103bp in bats and 1118 bp in rodents (Figure 2a). The phylogenetic analyses of the resulting fragments (identified by GARD) confirmed the recombination in both bats and rodents. Specifically, we found that the HERC-like 5’-fragment clustered with the HERC5 gene (clade supported by a significant bootstrap), while the 3’-fragment grouped with the HERC6 gene (Figure 2b, 2c). Taken together, our findings reveal that a similar ancient mechanism has independently led to the fixation of a HERC-like gene, which in fact is a chimeric HERC5/6 gene containing the HERC5 RLD domain and the HERC6 spacer and HECT domains in bats and rodents.

**Figure 2.**
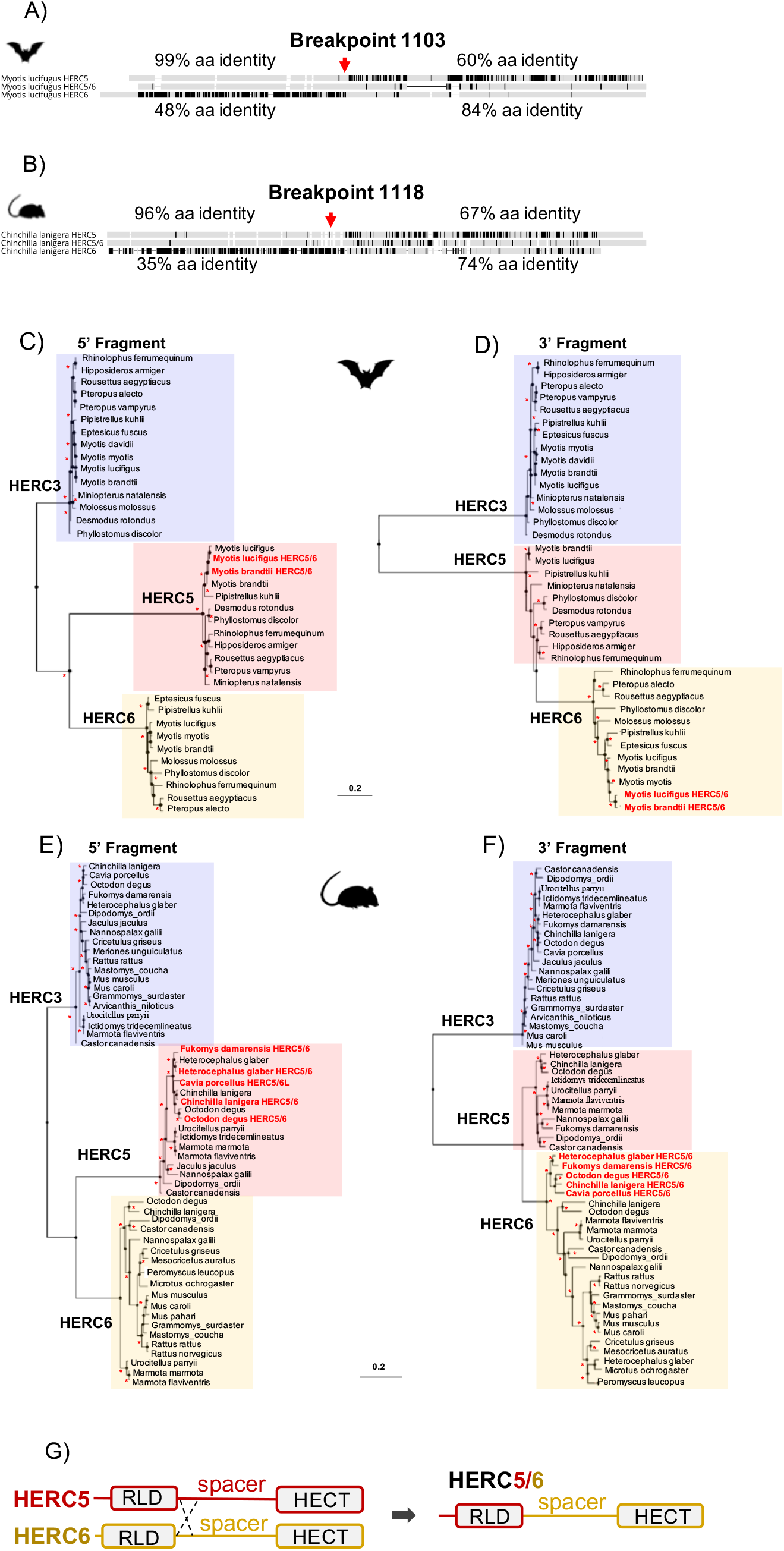
Independent duplication of a chimeric HERC5/6 gene through recombination in bats and rodents. (A, B) Alignment of the protein sequence of HERC5, HERC5/6, and HERC6 genes from bats and rodents, respectively. Additional sequence alignments are shown in Supplementary Figure 4. The percentages of pairwise amino acid identities between the N-terminals of HERC5/6 and HERC5 or HERC6, as well as the C-terminals of HERC5/6 and HERC5 or HERC6 are indicated. The significant recombination breakpoints (red arrows, p-value <0.05) assigned by the GARD program are shown for the rodent and bat HERC5/6 gene. (C-F) Maximum likelihood phylogenetic tree generated with the 5’ (at the left) and 3’ (at the right) of the HERC5, HERC6, HERC5/6, and HERC3 (as an outgroup) nucleotide gene alignment based on GARD recombination results (corresponding to the breakpoint 1103 in *Myotis lucifigus* in bats, and 1118 in *Chinchilla lanigera* in rodents). The duplicated chimeric HERC5/6 genes are shown in red. Asterisks indicate bootstrap values greater than 80%. The scale bar represents genetic variation of a 0.2 (20%) for the length of the scale. (G) A linear representation of HERC5 and HERC6 structures showing a chromosomal crossover between the 5’ regions, upstream of the spacer region. This mechanism has led to a duplicated recombined HERC5/6 gene containing the HERC5 RLD domain and HERC6 spacer region and HECT domain.

Moreover, by analyzing the phylogenetic trees of HERCs in bats and rodents (Figure 2b), we found that the 5’ fragment of HERC5/6 was genetically closer to HERC5 from the same species. This was not the case for the 3’ end, where all 3’ fragments of the HERC5/6 genes significantly grouped together. Combined with several GARD analyses, this supports that recent recombinations further occurred between the RLD domains of HERC5 and HERC5/6 genes.

### HERC6, but not HERC5, has been under strong positive selection during mammalian evolution

We next investigated whether HERC5 and HERC6 have been under selective pressure in mammals. Our results revealed some signatures of positive selection in artiodactyl, primate and bat HERC5 (p-values < 10^−3^), but none in rodent and carnivore species (p-value > 0.5) (Table 1). At the codon level, the signal was overall very weak, with less than three positively selected codons assigned per order (posterior probability threshold fixed at 0.95, and p-value < 0.05) (Table 2). In primates, two significant positively selected sites were identified in the RLD domain (Figure 3). Similarly, less than two codons evolved under positive selection in bats and artiodactyls (except for the MEME method which identified multiple positively selected sites in the artiodactyls), and none of the codons were common between methods (Table 2). Interestingly, the separate analysis of cetacean species revealed stronger signatures of positive selection at the gene (p-value = 6.10^−6^) and the codon levels (five sites identified by at least two methods), suggesting a lineage-specific adaptation.

**Table 1.**
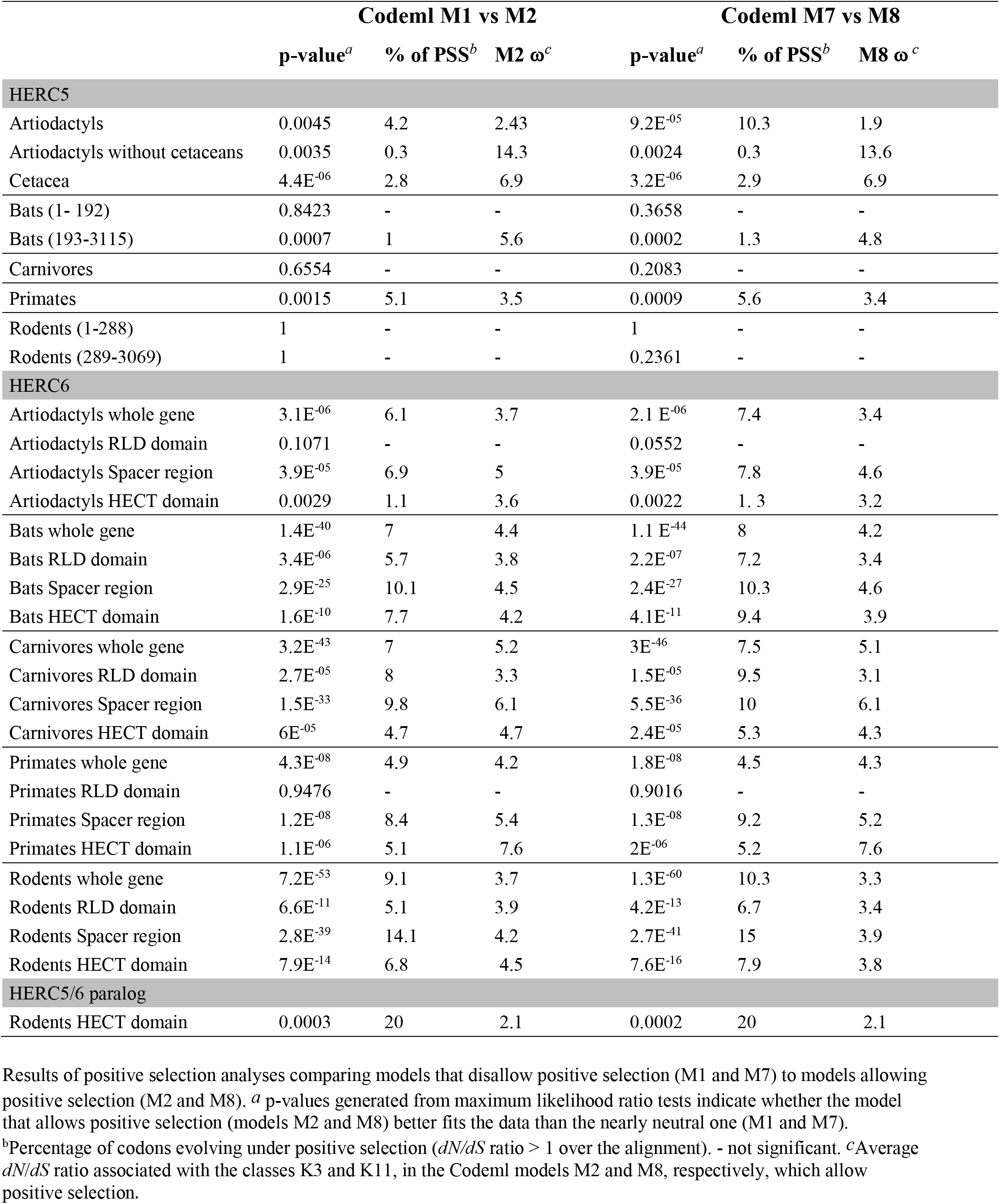
Positive selection analyses of mammalian HERC5 and HERC6 genes.

**Table 2.** Positively selected codons in mammalian HERC5, HERC6 and HERC5/6 genes.

|  | MEME | FUBAR | Codeml M2 | Codeml M8 |
| --- | --- | --- | --- | --- |
| **HERC5** |  |  |  |  |
| Artiodactyls | 6, 42, 63, 263, 408, 539, 565, 672, 696, 745, 992 | 39 | - | 189 |
| Artiodactyls without Cetacea | - | 39, 42, **552**, 555, 1034 | - | **552** |
| Cetacea | **6**, 664, 666, **683**, **712** | **6**, 175, 347, **663**, **676**, **683**, **712**, 732, **782** | **663**, **676** | **6**, 662, **663**, **676**, **712**, **782** |
| Bats (193-2913bp) | 22, 39, | 550 | - | 654 |
| Carnivores | 89, 90, 91, 93, 250, 265, 298, 323, 568, 573 | 189, 401, 439, 840 | - | - |
| Primates | - | **12**, **19**, **43**, 233, 296, **480**, **483** | **12**, **19** | **12**, **19** |
| Rodents (289-2773bp) | 59, 76, 87, 98, **191**, 363, 462, 658, 870 | **191** | - | - |
| **HERC6** |  |  |  |  |
| Artiodactyls | 56, 62, 112, **116**, 164, 167, 568, 661, 708, 723, 797 | 15, **116**, 599, 603, 655, 886 |  |  |
| Bats | **6**, 57, 58, 163, 382, 396, **549**, 552, 597, 600, **603**, **608**, 633, **646**, 650, **663**, 665, 671, **690**, **704**, **706**, 716, 814, 910, **911**, 995, 1014, 1015 | **6**, **549**, **603**, **608**, **642**, **646**, **647**, **648**, **663**, **690**, **704**, **706**, **911** | **6**, **18**, **74**, **454**, **546**, **549**, **591**, **595**, **603**, **608**, **642**, **646**, **647**, **648**, **650**, **651**, **663**, **690**, **706**, **911**, **964** | **6**, **18**, **74**, **454**, **471**, **546**, **549**, **591**, **595**, **603**, **608**, **642**, **643**, **646**, **647**, **648**, **650**, **651**, **663**, **690**, **706**, **911**, **964**, **974** |
| Carnivores | 30, 112, **166**, **216**, 225, **398**, **447**, 515, 517, **550**, 574, **593**, **596**, **600**, **602**, **603**, **659**, **666**, **669**, 753, 794, 846, 1023 | **166**, **216**, 397, **398**, **447**, 549, **550**, **593**, 595, **600**, **602**, **603**, **659**, **666**, **669**, 713, 968 | **9**, **23**, **63**, **70**, **166**, **216**, 395, **396**, **517**, **547**, **548**, **591**, **593**, **594**, **596**, **598**, **600**, **601**, **657**, **699**, **711**, **917**, **966** | **9**, **23**, **63**, **70**, **166**, **216**, **396**, **517**, **547**, **548**, **591**, **593**, **594**, **596**, **598**, **600**, **601**, **657**, **664**, **667**, **699**, **711**, **917**, **966** |
| Primates | 401, 472, 556, 647, 759, **828**, 1025 | 76, **597**, 631, 651, 700, **828**, 950, 989, 1021 | **597**, **641**, **710**, **826** | 370, 596, **597**, **641**, 649, 698, **710**, **826**, 987, 1020 |
| Rodents | **10**, **11**, **16**, 59, 66, 123, 209, 268, 275, **317**, 328, 366, 450, 474, **546**, **548**, **550**, 554, **602**, **606**, 609, **611**, **638**, **642**, **643**, 644, 649, 677, **686**, **688**, 697, 700, 743, 756, 797, 838, 842, 872 | **10**, **11**, **16**, **287**, **546**, **548**, **550**, **551**, **557**, **593**, **594**, **597**, **611**, **642**, **643**, **688** | **10**, **16**, **22**, **125**, **166**, **287**, **317**, **513**, **517**, **547**, **550**, **551**, **552**, **553**, **557**, **593**, **594**, **597**, **601**, **602**, **603**, **605**, **606**, **610**, **611**, **613**, **614**, **635**, **638**, **642**, **643**, **645**, **656**, **657**, **686**, **688**, **901**, **952** | **10**, **16**, **22**, **125**, **166**, **219**, **287**, **317**, **513**, **517**, **547**, **550**, **551**, **552**, **553**, **557**, **593**, **594**, **597**, **601**, **602**, **602**, **605**, **606**, **610**, **611**, **613**, **614**, **635**, **638**, **642**, **643**, **645**, **656**, **657**, **686**, **688**, **901**, **952** |
| **HERC5-6 paralogs** |  |  |  |  |
| Rodents | 69, 138, 261, **409**, **467**, 774, 926 | 29, 120, 361, **467**, 506, 507, **518**, 556, **566**, **660** | **467** | 22, **409**, **467**, **518**, **566**, **660** |

Results from site-specific positive selection analyses, with a posterior probability (PP) of BEB >0.95 for the models M2 and M8 from Codeml, >0.9 for FUBAR and a p-value <0.05 for MEME. Codons in bold are those that were assigned by at least two different methods. Codon numbering is based on HERC5 sequences from *Bos taurus*, *Tursiops truncatus*, *Phyllostomus discolor*, *Felis catus*, *Homo sapiens* and *Dipodomys ordii*, HERC6 sequences from *Bos taurus*, *Rhinolophus ferrumequinum*, *Felis catus*, *Homo sapiens* and *Mus musculus*, HERC5/6 sequence from *Fukomys damarensis*.

**Figure 3.**
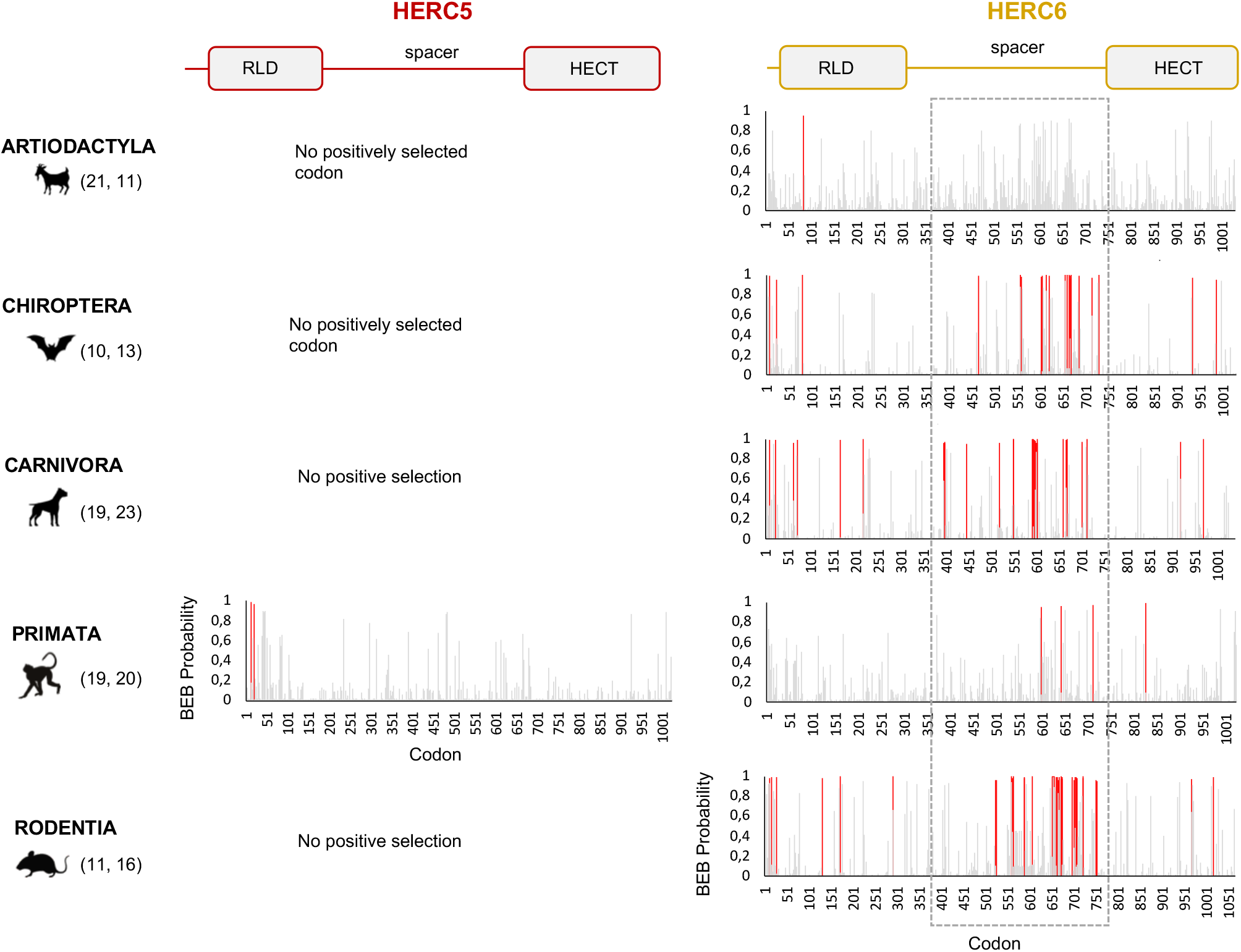
HERC6, and not HERC5, has experienced strong and mammalian-wide positive selection. Graphic panels represent the posterior probabilities of positive selection (Bayes empirical Bayes, BEB) (y axis) in the M2 Codeml model (allowing for positive selection, ω>1) for each codon (x axis) in HERC5 (left) and HERC6 (right) alignments. Red bars indicate the sites identified by both models, M2 and M8, with a BEB posterior probability greater than 0.95. Numbers in brackets are total species analyzed in each mammalian order for each gene. Site numbering is based on HERC5 protein sequences from *Homo sapiens*, HERC6 sequences from *Bos taurus, Rhinolophus ferrumequinum, Felis catus, Homo sapiens* and *Mus musculus.* Above is a linear representation of HERC5 and HERC6 showing the structural domains, the RLD, spacer region and HECT domains.

In contrast, we found very strong signatures of ancient and recurrent positive selection in HERC6, at both the gene and the codon levels. All five mammalian orders exhibited a significant excess of non-synonymous rate along the HERC6 coding sequences, specifically in bats, carnivores and rodents (p-value < 10^−43^ in bats, carnivores and rodents; and p-value < 10^−6^ in artiodactyls and primates). Positive selection was observed in each of the three domains of HERC6 – the RLD domain, the spacer region and the HECT domain – in bat, carnivore and rodent species, while many of the signatures were concentrated in the spacer region and the HECT domains in primates and artiodactyls (Table 1, Figure 3). Remarkably, the fastest-evolving codons mapped into the HERC6 spacer region of all five mammalian orders (p-value < 10^−26^ in bats, carnivores and rodents, p-value < 10^−5^ in artiodactyls and primates). More than 14 sites were identified by at least two methods in bats, carnivores and rodents, thereby constituting a hotspot of highly variable sites in mammalian HERC6 (Table 2 and Figure 3). In line with this finding, alignment of the spacer region revealed that it is an extremely divergent domain, characterized by multiple amino acid changes and indels between and within groups, in particular in rodents and bats (Figure 4).

**Figure 4.**
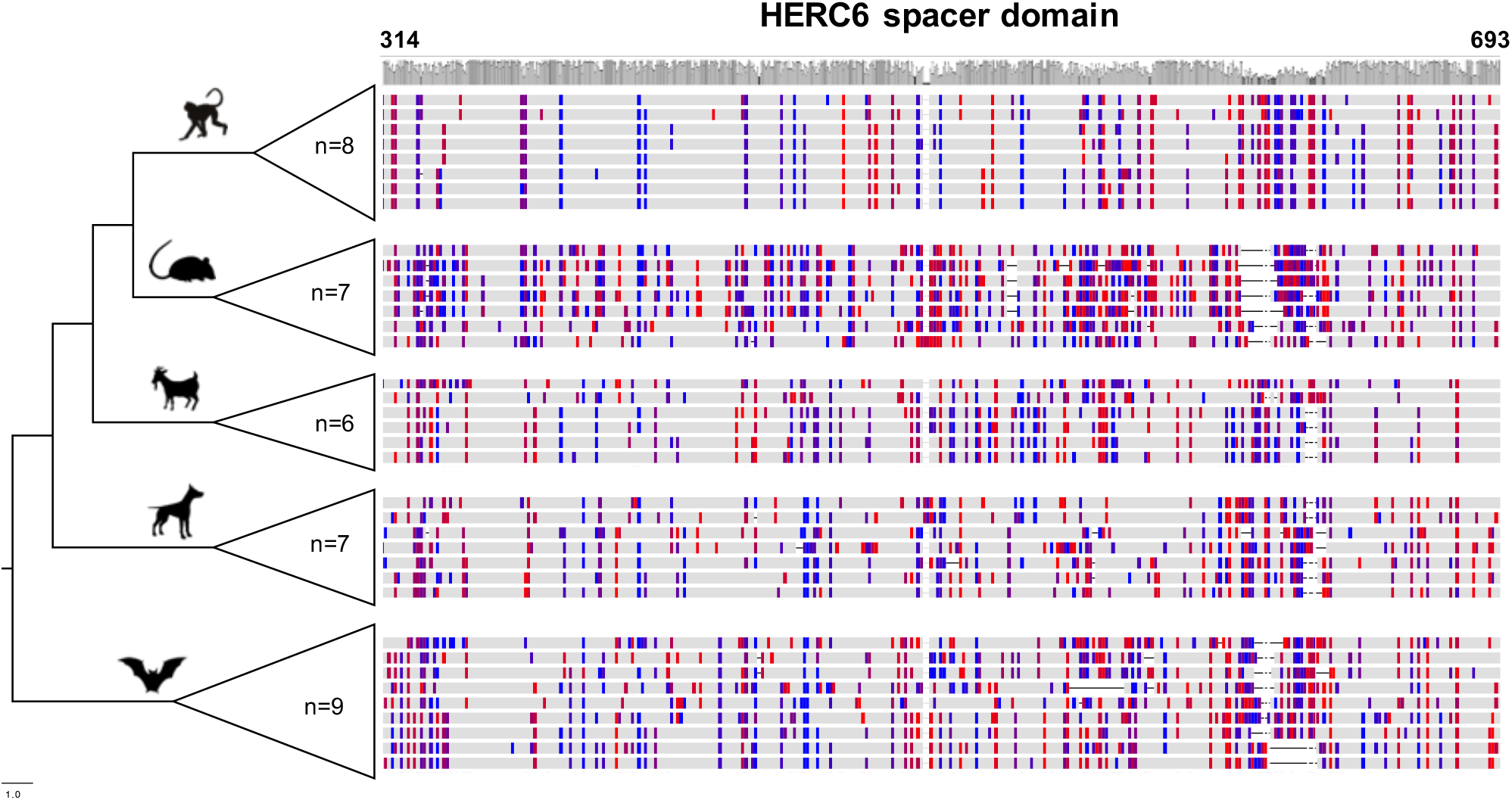
Rapid evolution of mammalian HERC6 spacer region is characterized by multiple amino acid changes and major indels. Multiple alignment and comparison of the HERC6 spacer region between and within mammalian orders. Left, cladogram with the number of sequences used for each clade (n=6 to 9). Right, colors indicate site variations between the sequences as compared to the consensus sequence with a threshold of 25% (Geneious, Biomatters; blue/red, hydrophilic/hydrophobic residues), while grey represents similarity with the consensus. The average pairwise percentage of identity is graphically represented above in gray. The codon numbers are based on human HERC6 sequence.

Therefore, although HERC5 presents low evidence of positive selection in mammals, HERC6 has experienced very strong adaptive evolution. Both genes showed differential evolutionary profiles across/between mammals, with lineage-specific and domain-specific adaptations.

### The rodent chimeric HERC5/6 paralog has evolved under strong positive selection

We then addressed whether the newly identified chimeric HERC5/6 gene, which contains the HERC5 RLD domain and the HERC6 spacer and HECT domains, has also experienced positive selection. As coding sequences from five rodent species were available, we assessed the inter-species evolutionary history of the chimeric HERC5/6 gene within this group. Of note, there were insufficient bat sequences/species to perform the corresponding analyses. The likelihood ratio tests revealed significant positive selection in rodent HERC5/6 (p-value<0.0003, Table 2). Importantly, all the positively selected codons mapped in the spacer region, and were concentrated between the amino acids 409 and 660 (Figure 3), suggesting that the spacer domain has been the target of strong positive selection as observed in the HERC6 protein.

### Rodent and bat HERC6 genes have been under stronger positive selection compared to other mammals

Finally, we tested whether positive selection has differentially impacted the evolution of HERC5 and HERC6 across mammals. We found that the chiropteran and rodent HERC6 genes have experienced intensive episodic positive selection compared to the other groups (Figure 5). In particular, five chiropteran lineages distributed along bat phylogeny (*Rhinolophus ferrumequinum, Pteropus* ancestral branch, *Phylllostomus discolor, Molossus molossus*, and *Pipistrellus kuhlii*) and five rodent branches (two ancestral branches of mouse related clade, *Cricetulus griseus, Mesocricetus auratus*, and *Urocitellus parryii*) have undergone a significant excess of amino acid changes, with a ratio ω > 3.8. Differential selective pressure in HERC5 was also evidenced across mammalian lineages, but to a lesser extent: only two branches were found under significant positive selection, in bats (*Hipposideros armiger*, ω=189) and carnivores (*Ailuoropoda melanoleuca*, ω=78).

**Figure 5.**
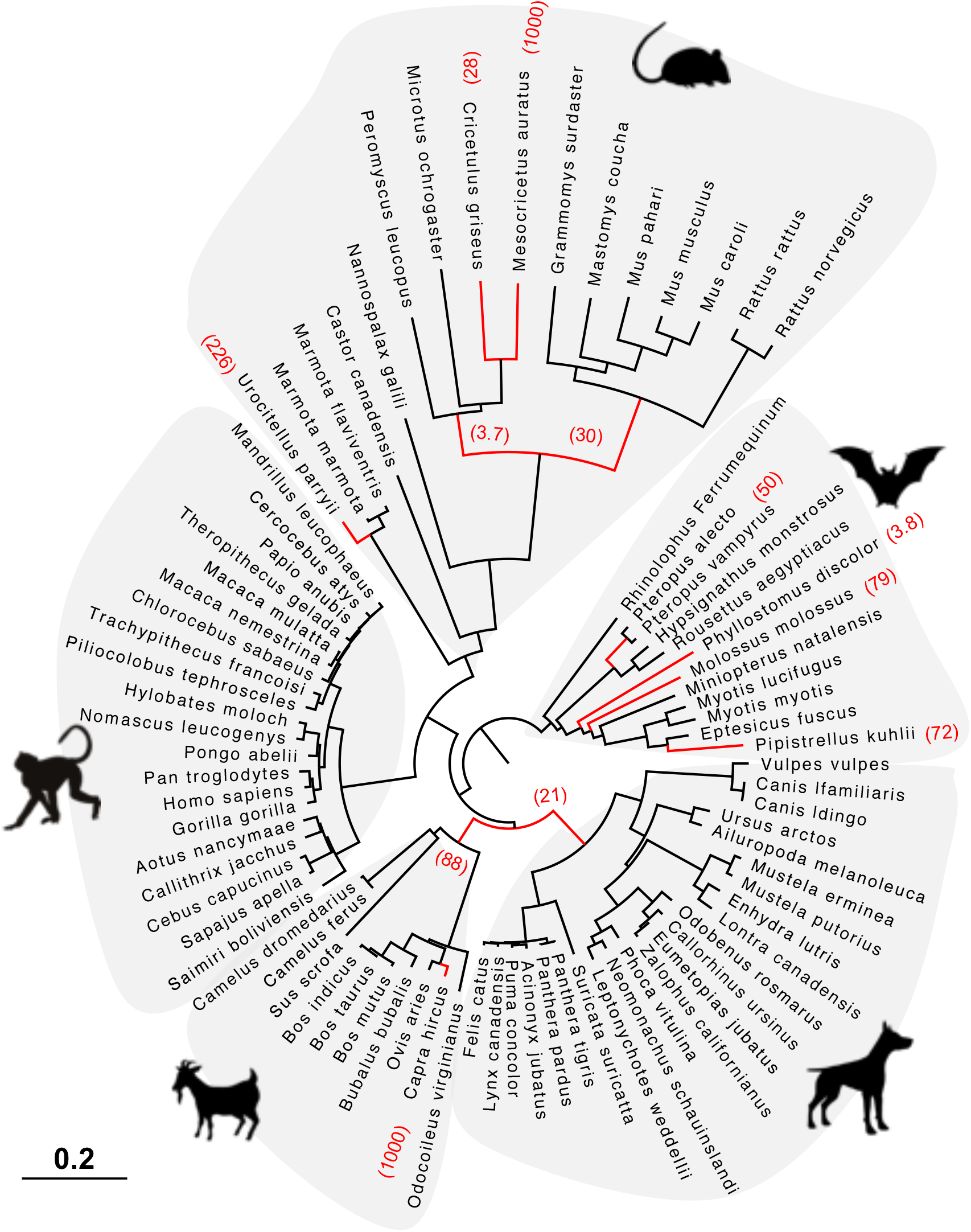
HERC6 has been under strong positive selection during rodent and bat evolution. Maximum likelihood phylogenetic tree of mammalian HERC6 gene showing the branches under significant positive selection (p-value < 0.05, in red) assigned by aBSREL from the HYPHY package. The numbers in brackets indicate the estimated values of the ω at the branch. The scale bar indicates the proportion of genetic variation.

## Discussion

Deciphering the evolutionary and functional diversification of the antiviral innate immunity in mammals is of primary importance to better understand modern viral pathogens, virus-host interfaces, and identify novel antiviral strategies. The functional significance of HERC5 and HERC6 is underlined by their ancient origin and conserved expression in vertebrates (54). However, their evolutionary history in mammals has remained unclear. Here, we have carried-out in-depth phylogenetic and genomic analyses to address how mammalian HERC5 and HERC6 have evolved over millions of years of divergence.

### Differential evolutionary fate of HERC5 across mammals

Although mammalian HERC5 was previously reported as a rapidly evolving gene (36), we only found strong evidences of recurrent positive selection in primates. In particular, two codons in primate HERC5 have rapidly evolved in the RLD domain, possibly reminiscent of pathogen-exerted pressure (3,55–57). In line with this, blade 1 of the primate RLD domain was recently reported to be an important functional region for HERC5 anti-HIV antiviral activity (54). Thus, retroviruses may have played a role in the diversification of HERC5 during primate evolution. Such patterns have been reported in many primate restriction factors, including BST2 (58–60), TRIM5 (10,61–63) and APOBEC3 (14,64–66). However, because the role of HERC5 is not limited to host defense against retroviruses, its evolution in primates may also reflect past selection against other viral pathogens such as influenza viruses and papillomaviruses.

In contrast, positive selection was solely evident at the gene level for artiodactyl and bat HERC5, and absent in rodents and carnivores. This pattern may be a result of lineage-specific selective drivers: differential viral exposure history, distinct mechanisms of viral antagonism, and/or may reflect overall pressure to maintain efficient cellular functions of HERC5. For example, apart from its antiviral role, some evidences suggest that HERC5 might be functionally involved in other pathways, such as spermatogenesis and cell cycle (22), as well as cancer (67). HERC5 may have thus evolved to maintain effective cellular functions rather than to escape viral antagonisms or to target viral pathogens in bats and artiodactyls. In mammals not exhibiting positive selection, viruses may have evolved indirect mechanisms of antagonism to counteract HERC5 function. In line with this, many viruses encode proteins that interfere with the ISGylation activity of HERC5, through direct interaction with the ISG15 protein (68–71). For example, the NS1B protein encoded by influenza virus antagonizes ISG15 conjugation through direct interaction (68).

### Accelerated evolution of HERC6 in mammals

HERC6 is the only protein from the small HERC family exhibiting such high levels of adaptive changes with an extremely divergent spacer region in all mammals, except artiodactyls. Such rapid evolution of HERC6, with accumulated mutations replacing the amino acids and multiple amino acid insertions/deletions, most likely mirrors pathogen-driven adaptations as a result of past evolutionary arms-races (3,55–57). This highlights a fundamental antiviral role for HERC6 in mammals. Previous functional evidences support that HERC6 is involved in mammalian antiviral immune response (28,34,35,68). However, available studies have only focused on HERC6 anti-retroviral activity and ISGylation function. For example, HERC6 has been shown to be an IFN-inducible E3-ligase of ISG15 conjugation in mouse (28,34,35). In contrast, human HERC6 lacks the ISGylation activity, but it potently inhibits the primate lentivirus SIVmac viral production (54).

Specifically, we identified the spacer region as a potential pathogen – HERC6 interface, involved in the recognition of viral proteins and/or being the target of viral antagonists. Up to now, most studies have been devoted to the functional characterization of the RLD and HECT domains of HERC proteins, as they belong to the well-characterized protein families, RCC1 and E3-ligases, respectively. In contrast, the structural characteristic of the spacer region has not been related to any known protein, which hinders its description and functional role. However, its propensity for amino acid insertions/deletions and accumulated non-synonymous mutations, including changes with strong chemicophysical differences highlights a strong evolutionary plasticity. Such a hotspot of variability in unstructured regions has been reported for several restriction factors, such as MX1, in which the highly variable L4 loop has led to differential virus-host interfaces and plasticity (72,73).

It is noteworthy that the RLD and HECT domains of HERC6 were also subjected to positive selection in bats, carnivores and rodents. These signatures may reflect different virus-host interfaces. This pattern was reported in the primate PKR protein, in which signatures of pathogen-driven selection are scattered along the protein as a result of interactions with multiple viral antagonists (74,75). However, it is also possible that all positively selected sites cluster in a same spatial region of the protein. A 3D structural analysis of the HERC6 protein would help assessing how the rapidly evolving sites are located in the protein, and allow determining whether HERC6 presents multiple or unique host-pathogen interface(s). Up to now, the 3D structure has only been solved for the HECT domain and RLD domain separately [e.g. (76,77)]. Therefore, how the spacer region connects the HECT and RLD domains in a 3D structural dimension is currently unknown. Further studies on small HERC protein structure would help better understanding how the high variability in HERC6 impacts its structure and function.

We found differential genetic profiles of HERC6 across mammalian orders. The strongest signal was found in bats, carnivores, and rodents, suggesting that different strength / intensity of selective pressures have shaped the evolution of HERC6 in mammals. This was confirmed by the branch-specific analyses, in which rodents and bats exhibit significant lineage-specific adaptive changes. Rodents and bats are the two most diverse mammalian orders, and host the highest viral richness among mammals (39). Both orders have thus been exposed to a greater viral diversity, compared to primates and artiodactyls. This may have increased the strength and extent of selective pressure exerted on the HERC6 protein.

### Unequal recombination has led to the duplication of a chimeric HERC5/6 in rodents and bats

Lineage-specific expansions of multigene families have shaped and complexified the mammalian antiviral repertoire over million years of evolution. Consequently, many unrecognized genes encoding for antiviral proteins are yet to be discovered. Here, we have identified duplications of a HERC paralog in the rodent Hystricognathi infra-order and the chiropteran *Myotis* genus, which probably occurred around 30 MYA (78,79) and 44 MYA (80–82), respectively.

Interestingly, these paralogs are chimeric HERC5/6 genes coding for the HERC5 RLD fused to the HERC6 spacer region and HECT domain. This suggests that an independent duplication has occurred through a similar mechanism in bats and rodents. Gene duplication can occur by different modes, mainly including unequal crossing over, retroposition or chromosomal duplication (83,84). In the former case, duplicated genes are physically linked in the chromosome, and can contain a fragment of a gene, a whole gene or several genes (83). In contrast, retroposition engenders a retrotranscribed complementary DNA incorporated into the genome, which generally lacks intronic regions and regulatory sequences (83). Given that the HERC5/6 are located in the canonical locus of HERC5 and HERC6 and contain the parental intronic regions, they have most likely resulted from a unequal crossing over between HERC5 and HERC6.

This hypothesis is supported by the recombination analyses, which detected a significant breakpoint upstream of the spacer region in both mammalian orders. The fact that the recombination occurred at the same genetic location can be explained by two different, but not necessarily mutually exclusive, hypotheses. First, the recombination event can only occur at this location because of genomic structural constraints (i.e. genomic homology between paralogs). Second, the HERC6 spacer region and HECT domains are required for functional HERC5/6 proteins. The best example of such tandem duplication with domain fusion is the lineage-specific expansion of the APOBEC3 family in mammals. Originating from an ancestral APOBEC3 gene, tandem duplications as well as retrocopying events have radically expanded the repertoire of mammalian APOBEC3 genes (14,20). In primates, several of the APOBEC3 genes have most likely resulted from the fusion of A3 domains, while the murine genome encodes a unique APOBEC3Z2-APOBEC3Z3 fused gene (85), highlighting lineage specific functional adaptations.

Such gene duplications accompanied with gene fusion are major genetic innovations that functionally diversify the antiviral arsenal (3,4,7). For example, the expansion of the APOBEC3 family has functionally diversified the antiviral activities and specificity of targeted viruses in primates (20,64,86). Given the antiviral role of HERC5 and the potential implication of HERC6 in antiviral immunity, we hypothesize that the HERC5/6 paralog provides a functional advantage against pathogenic viruses. This is supported by both the extremely rapid episodic evolution of HERC6 in rodents and bats, and the signatures of positive selection in rodent HERC5/6. The fused HERC5/6 gene may have evolved combined functional features of HERC5 and HERC6. Alternatively, it may have retained the functional implication of the HERC5 RLD domain, but has functionally diverged from HERC6.

This latter hypothesis is more likely as we found that the RLD domain of HERC5 and of HERC5/6 are highly similar and cluster together within species (more than 95% pairwise amino acid identity). This pattern may reflect ongoing gene conversions between the RLD domains of HERC5 and HERC5/6, thereby maintaining a N-terminal similar to the parental HERC5 protein.

Whether the independent fixation of the HERC5/6 paralog in rodents and bats is a functional/phenotypic convergent evolutionary event has to be investigated. Moreover, it raises the question why both lineages have undergone such genetic innovation. While this duplication could be hazardous, it is possible that the rodent Hystricognathi infra-order and the bat *Myotis* genus share a common selective pressure, such as a viral pathogen family. Other forces such as ecological and/or environmental factors may have played a role, such as life-history traits or biodiversity changes.

### Perspectives

Studying the genetic adaptations of host innate immunity can provide insights into the evolutionary and functional determinants of host antiviral response. This approach has been a powerful tool for assessing the functional diversification of virus - host interfaces in many systems [e.g. (61,64,87,88)]. In this present study, we identified HERC6 and HERC5/6 as potential unrecognized restriction factors in mammals. Further functional investigations are now required to (i) decipher the antiviral function of HERC6 and HERC5/6, (ii) determine how the accumulated variability in the spacer domain may impact their structure, function and stability, (iii) unravel HERC6 binding interface with potential viral antagonists or targeted viral proteins, and (iv) determine on the other side which pathogens are targeted by those proteins. Moreover, deciphering whether and how the HERC5/6 paralogs afford a selective advantage against pathogens in rodent and bat lineages is of main interest in virology and immunology fields, as both orders are important reservoirs of zoonotic pathogens. Finally, studying the functional implications of HERC6 adaptation in mammals will not only allow to better understand how pathogens have shaped host immunity, but will provide important insights into the overlooked role of small HERC proteins in mammalian antiviral response.

Altogether, our results represent avenues for future functional studies of importance in mammalian innate immunity.

## Acknowledgements

We are particularly grateful to Adil El Filali (UMR5558) for his help on genomic analyses, and Laurent Duret (UMR5558) for his comments on the study. We thank Andrea Cimarelli, head of the Host-Pathogen Interaction during Lentiviral Infection team at the CIRI Lyon, as well as the members of the “Intercontinental Retrovirus Zoomposium” for helpful discussion. We also thank all the contributors of the LBBE bioinformatic server and the publicly available genome sequences.

This work is funded by the ANR LABEX ECOFECT (ANR-11-LABX-0048 of the Université de Lyon, within the program Investissements d’Avenir [ANR-11-IDEX-0007] operated by the French National Research Agency). Lucie Etienne is further supported by the CNRS and by grants from amfAR (Mathilde Krim Phase II Fellowship no. 109140-58-RKHF), the Fondation pour la Recherche Médicale (FRM Projet Innovant no. ING20160435028), the FINOVI (“recently settled scientist” grant), the ANRS (no. ECTZ19143 and ECTZ118944), and a JORISS incubating grant. Dominique Pontier is supported by the CNRS, the European Regional Development Fund (ERDF), and the ANR EBOFAC.

## Author contributions

SJ carried-out the analyses of the data. SJ, LE and DP performed the investigations. SJ, LE and DP wrote the original draft of the paper. LE and DP acquired the funding, and coordinated the study. All authors reviewed and edited the manuscript.

## Supplementary information

**Supplementary Figure 1.**
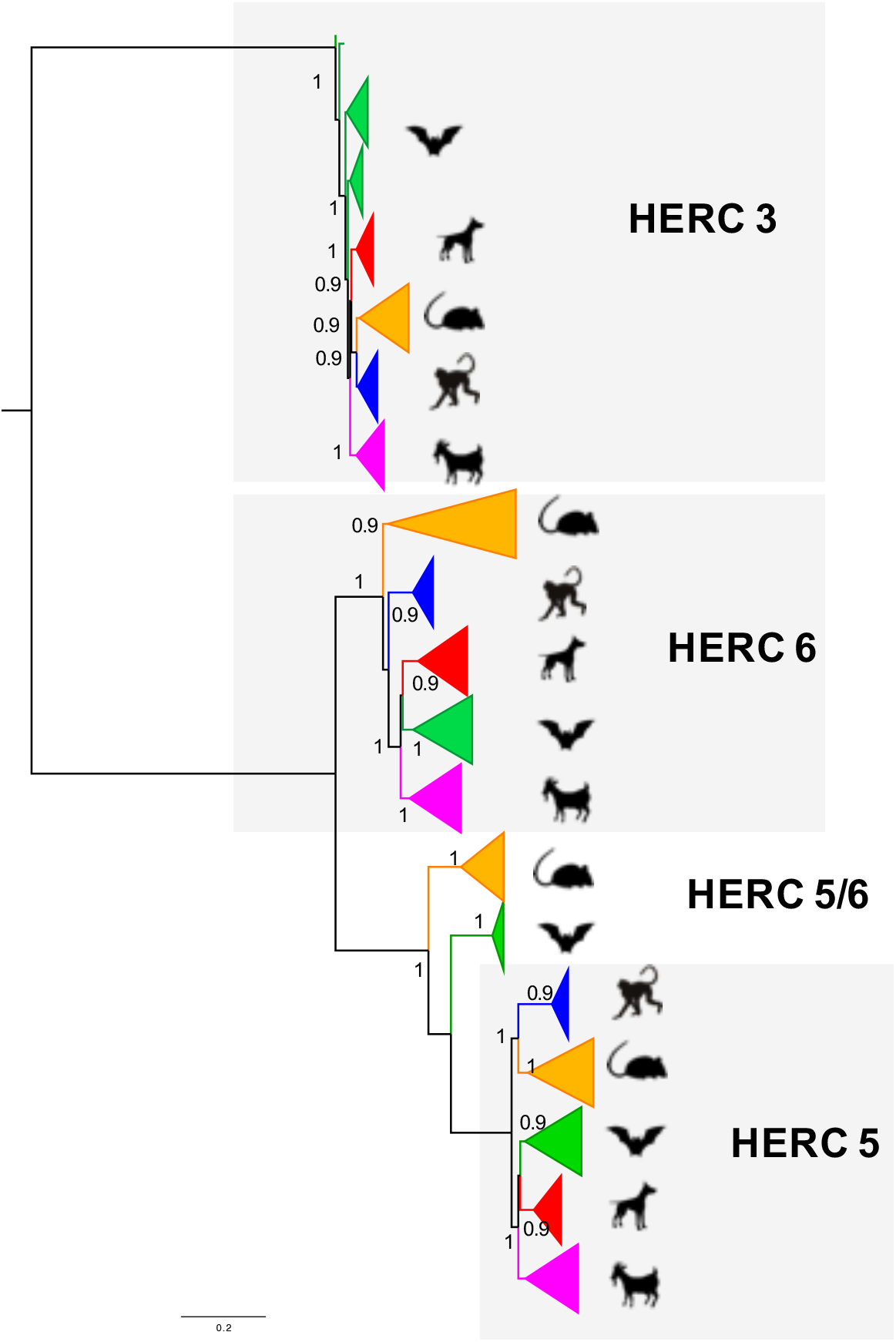
Maximum likelihood phylogenetic tree generated with the whole coding sequences of HERC5, HERC6, and HERC3 nucleotide alignment from artiodactyl, carnivore, rodent, bat and primate species. Asterisks indicate bootstrap values greater than 80%. The scale bar represents the proportion of genetic variation (0.2 for the scale), and is indicated at the bottom. Sequences are collapsed in each order for better readability.

**Supplementary Figure 2.**
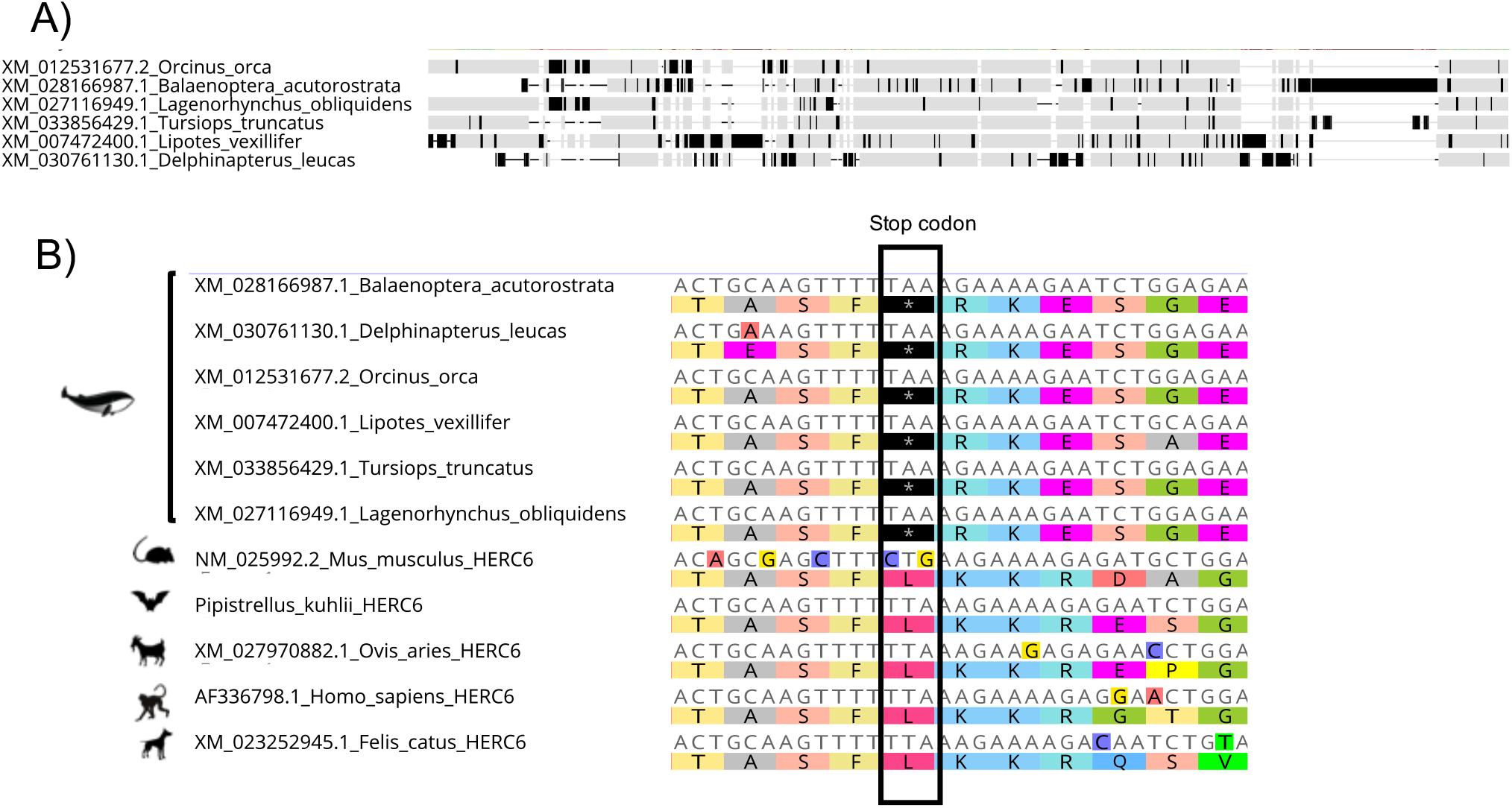
Pseudogenization of cetacean HERC6. A) Multiple amino acid alignment of six cetacean HERC6 sequences showing multiple substitutions, insertions and deletions. B) Multiple nucleotide alignment with corresponding amino acids of cetacean, rodent, bat, ruminant, primate and carnivore HERC6, highlighting a conserved stop codon in the cetacean species (codon 174 in *Balaenoptera acutorostrata*). Nucleotide and amino acid sequences are shown using Geneious.

**Supplementary Figure 3.**
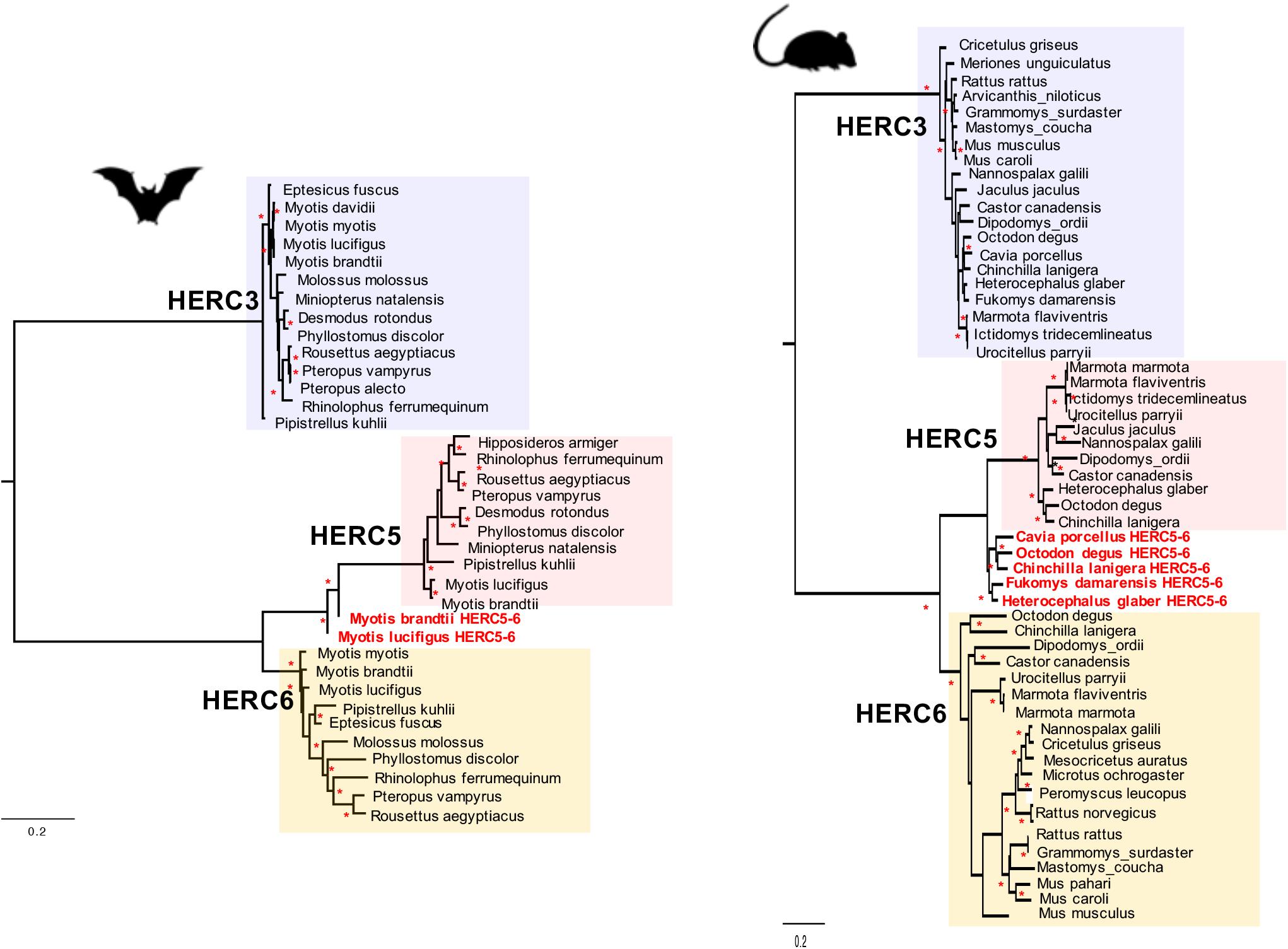
Maximum likelihood phylogenetic tree generated with the whole coding sequences of HERC5, HERC6, HERC5/6, and HERC3 nucleotide alignment in bats (left) and rodents (right). The chimeric duplicated HERC5/6 genes are shown in red. Asterisks indicate bootstrap values greater than 80%. The scale bar at 0.2 is indicated below.

**Supplementary Figure 4.**
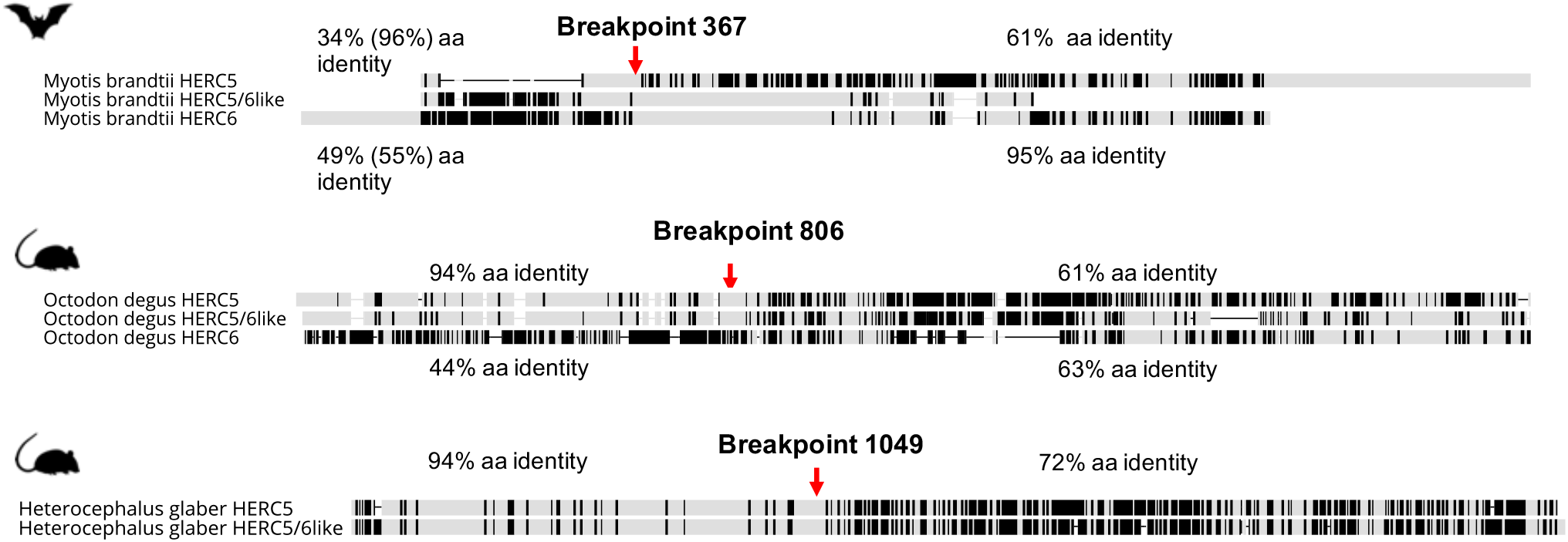
Alignment of the protein sequence of HERC5, HERC5/6, and HERC6 from bats and rodents. The percentages of pairwise amino acid identities between the N-terminals of HERC5/6 and HERC5 or HERC6, as well as the C-terminals of HERC5/6 and HERC5 or HERC6 are indicated. The significant recombination breakpoints (red arrows, p-value <0.05) assigned by GARD program are shown for bat and rodent HERC5/6 gene. Because the coding sequence of HERC6 gene from *Heterocephalus glaber* was incomplete with many missing data, it was not included in the figure. Likewise, some portions of the N-terminal of HERC5, as well as the C-terminals, from HERC5/6 and HERC6 are missing in the protein alignment of the chiropteran species, *Myotis brandtii* (top).

**Supplementary Table 1.**
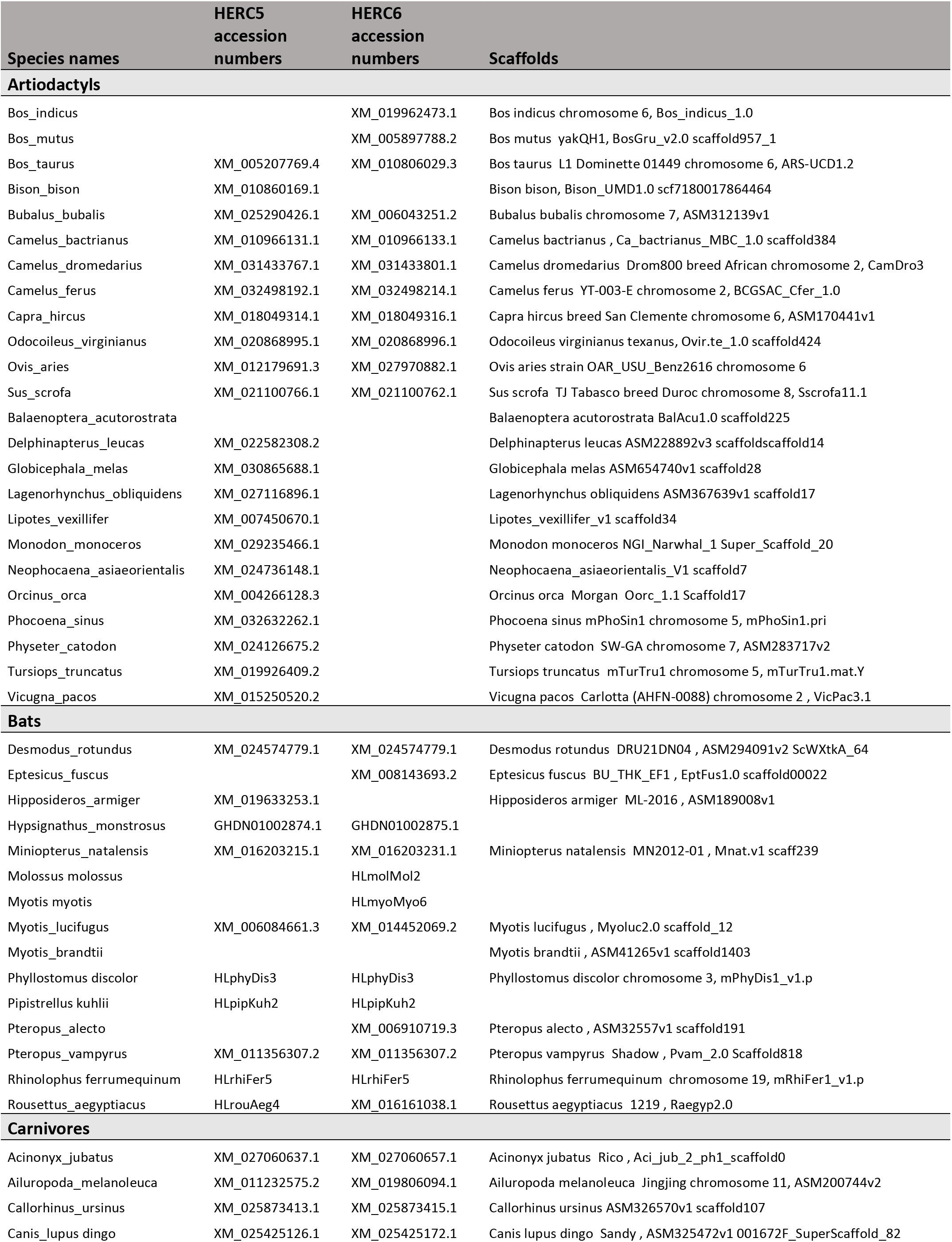
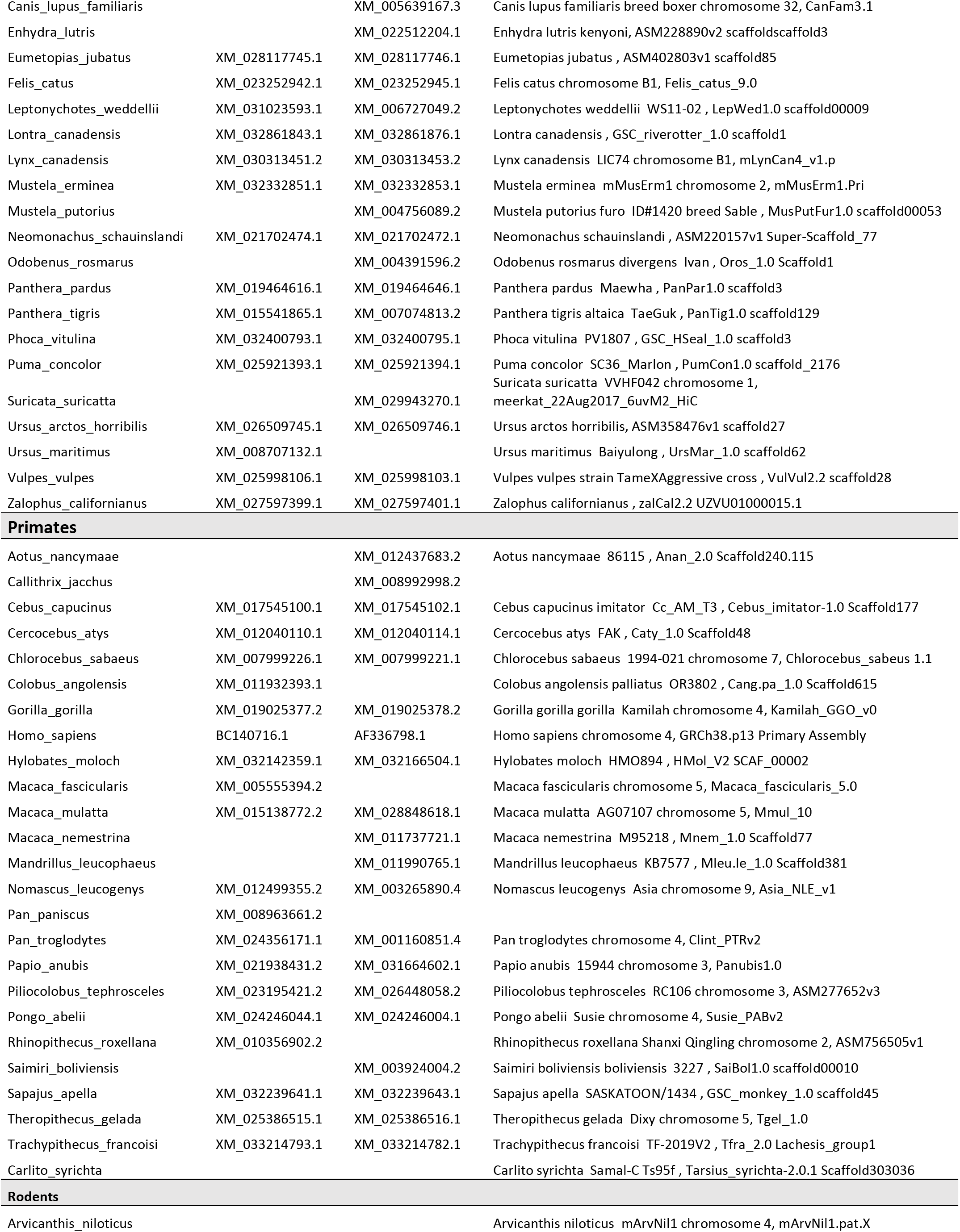
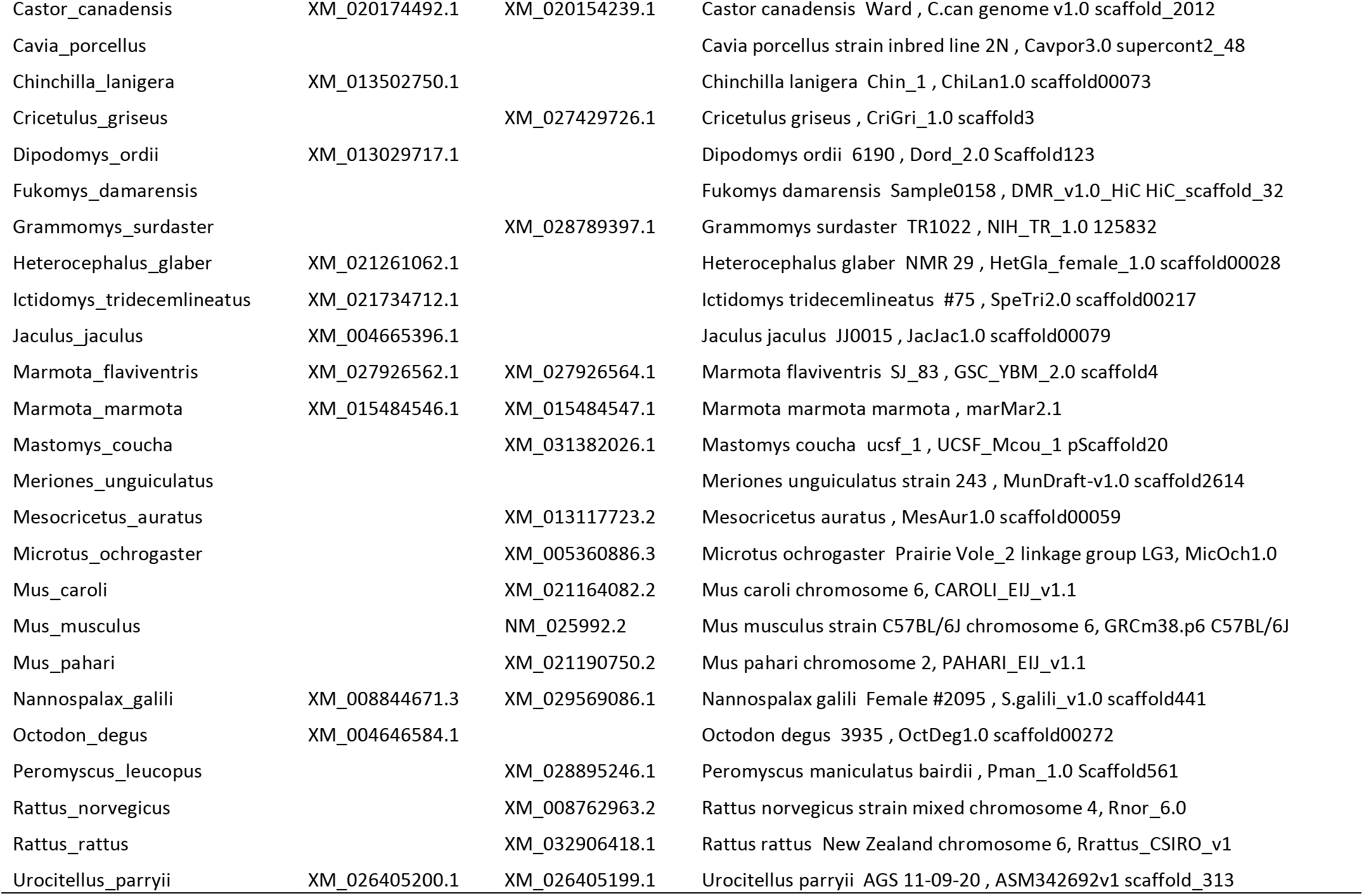
Information on publicly available datasets analyzed in this study. Accession numbers are available in NCBI (https://www.ncbi.nlm.nih.gov/).

